# Feature selection for the accurate prediction of septic and cardiogenic shock ICU mortality in the acute phase

**DOI:** 10.1101/337261

**Authors:** Alexander Aushev, Vicent Ribas Ripoll, Alfredo Vellido, Federico Aletti, Bernardo Bollen Pinto, Antoine Herpain, Emiel Hendrik Post, Eduardo Romay Medina, Ricard Ferrer, Giuseppe Baselli, Karim Bendjelid

## Abstract

Circulatory shock is a life-threatening disease that accounts for around one-third of all admissions to intensive care units (ICU). It requires immediate treatment, which is why the development of tools for planning therapeutic interventions is required to deal with shock in the critical care environment. In this study, the ShockOmics European project original database is used to extract attributes capable of predicting mortality due to shock in the ICU. Missing data imputation techniques and machine learning models were used, followed by feature selection from different data subsets. Selected features were later used to build Bayesian Networks, revealing causal relationships between features and ICU outcome. The main result is a subset of predictive features that includes well-known indicators such as the SOFA and APACHE II scores, but also less commonly considered ones related to cardiovascular function assessed through echocardiograpy or shock treatment with pressors. Importantly, certain selected features are shown to be most predictive at certain time-steps. This means that, as shock progresses, different attributes could be prioritized. Clinical traits obtained at 24h. from ICU admission are shown to accurately predict cardiogenic and septic shock mortality, suggesting that relevant life-saving decisions could be made shortly after ICU admission.

## Introduction

Shock (or circulatory shock) is a life-threatening medical condition that requires immediate treatment. It is prevalent in the Intensive Care Unit (ICU) and a major concern in Critical Care in general. This condition occurs when the organs and tissues of the body do not receive enough blood and, as a result, cells see their oxygen and nutrients supply restricted so that organs become damaged. Hypotension, tissue hypoperfusion and hyperlactatemia are amongst the most common symptoms [28]. The outcome of an individual case depends on the stage of shock, the underlying condition and the general medical state of the patient [28].

Four types of shock are commonly defined: hypovolaemic shock (e.g. hemorrhagic shock), cardiogenic shock, distributive shock (e.g. septic shock) and obstructive shock. The mortality rate of the condition remains very high and depends on its type. It is quantified at 30% for septic shock (according to the new definition of such condition [29]) and between 35.9% and 64.7% for in-hospital mortality in patients with cardiogenic shock (depending on the stage, [30]). Usually, shock is treated in the ICU, where it is carefully monitored. As a result, large amounts of data are produced for each patient.

Machine Learning (ML) techniques can help find a list of attributes to predict the outcome of shock patients from clinical data (i.e. data routinely monitored in the ICU). However, there is a large amount of attributes that characterize each patient, ranging from base pathology information, to lab procedures or hemodynamic data and treatment, to name a few. As a result, data are likely to be very high dimensional and, therefore, feature selection (FS) methods are often required to reduce data dimensionality and its associated variability. The combination of statistics and ML may provide an appropriate framework to retrieve new knowledge from data, as well as causal relationships between different features from high-dimensional datasets [31–33].

This study analyzes the clinical database gathered within the *ShockOmics* research project. The main objective of this project was to find ‘*omics*’ signatures of shock and it included two types of shock: septic and cardiogenic. Septic shock is the most common, and it is caused by infection. The second most frequent form of shock is the cardiogenic shock and it is a result of a heart failure. The multiscale nature of the *ShockOmics* database allows to search for attributes that are related to mortality induced by these two types of shock.

When it comes to the detection of circulatory shock, physicians and therapists largely depend on a combination of clinical, hemodynamic and biochemical signs. For a treatment to be chosen, prompt identification of shock is necessary. Appropriate treatment is based on a good understanding of the physiological mechanisms behind the condition. The management of shock is challenging and patient survival is highly dependent on the timely administration of the appropriate treatment. Normally, it requires the administration of vasoactive drugs and fluid resuscitation. This treatment is usually given to counteract the other conditions that go along with shock (e.g. hypotension/hemodynamic instability, inflammation and multiple organ failure (MOF)). The fact that they have similar symptoms makes addressing the causes of shock very difficult. As a result, decision making at the onset of the condition is not a trivial task and, therefore, it is important to design fast, reliable and interpretable tools to plan therapeutic interventions to prevent mortality, or any irreversible consequences caused by shock.

The study of the pathophysiology of shock and its management has been an active area of research in ML applications for Critical Care. For example, fuzzy decision support systems (DDS) for the management of post-surgical cardiac intensive care patients have been described in [4], while the issue of rule generation for septic shock patients has been addressed in [5] and [6], the former together with an artificial neural network (ANN). The expert system described in [8] is more closely related to the general management of sepsis and it was designed for the detection of pathogens and the prescription of antibiotics during the first hours of evolution. Ross and co-workers [9] later derived a system of ordinary differential equations together with an ANN model of inflammation and septic shock. Other studies have deployed ANNs for the study of sepsis. They include [7], who presented a clinical study examining SIRS and Multiple organ dysfunction syndrome (MODS) in the ICU after cardiac and thoracic surgery. More recently, a state-of-the-art application of a Deep Learning (DL) technique, namely Deep Reinforcement Learning (DRL) has been proposed by Raghu and co-workers [10] for the definition of continuous-space models for the treatment of sepsis, in a twist that goes beyond the more traditional development-and-use of discriminative classifiers. Support vector machine (SVM) models have also been used for the prediction of sepsis. Kim and co-workers [11] applied them to study the occurrence of sepsis in post-operative patients. Wang *et al*. [12] went a step further to build a DSS for the diagnosis of sepsis.

ML methods have also been used with varying success for the more specific problem of the prediction of mortality caused by sepsis. A diagnostic system for septic shock based on ANNs (Radial Basis Functions -RBF- and supervised Growing Neural Gas) was presented in [14]. Also in this area, Brause and colleagues [15] applied an evolutionary algorithm to an RBF network (the MEDAN Project) to obtain, over a retrospective dataset, a set of predictive attributes for assessing mortality for Abdominal Sepsis. Relevance Vector Machines (RVM), SVM variants with embedded feature selection, were used in [19], while Bayesian Networks were used in [16] and [17]. Finally, kernel methods were used in [18].

The work reported in this paper attempts to identify clinical traits that can be used as predictors of mortality in patients with cardiogenic or septic shock. Missing data imputation techniques and ML models are used, followed by different approaches to FS from different data subsets. Selected features are later used to build causal Bayesian Networks (CBN), in order to reveal potential causal relationships between features and ICU outcome.

The remaining of the paper is structured as follows. The section Materials and methods describes the *ShockOmics* dataset, as well as the FS, the ML and structure learning methods used in the reported experiments. The section Experimental Results presents the results for the FS experiments (obtained feature subsets and their classification performances) and causal discovery experiments (CBN structures). The experimental results are discussed in the section Discussion. The last section of the paper draws some conclusions about the investigated clinical problem.

## Materials and methods

The ML pipeline presented in this paper is divided into two main experimental phases: FS and causal discovery.

The FS experiments deal with the problem of finding the most promising attributes (clinical traits) for the prediction of mortality related to shock. To do this, three subsets of the *ShockOmics* data were analyzed: *T1*, *T1+T2* and *Full* (described below). The goal of the experiments was to analyze different feature sets resulting from the FS procedure, and to measure their performance using ML models. The decision of starting the analyses with three different data subsets of the *ShockOmics* database was made due to the importance of obtaining a stable subset of features that behave consistently at different time-steps and that are clinically relevant. For this reason, each of the datasets targeted different features: some of them because of their low amount of missing values, and some of them because they were important at different times. There is an assumption behind the experiments, namely that the feature sets with the best performances are also the most promising for mortality prediction.

The evaluated feature sets were obtained by applying four different FS techniques to the aforementioned datasets: Univariate FS based on ANOVA F-value (*UFS*); recursive feature elimination with Support Vector Classification (*RFE*); two stage FS (*UFS+RFE*) and Random Forest (*RF*) FS. The additional fifth method was applied by averaging stability scores and aggregating features from the four previous FS techniques results. Missing values were imputed using RF imputation separately in the training and test sets. The FS techniques were performed only on the training set. The identification of the promising feature sets was based on both performance and stability scores.

Finding a subset of attributes that are predictive of mortality due to shock may not suffice in a clinical setting. The causal discovery experiments reported next allow us to go one step further and analyze the causal relationships between the data features that were selected in the previous phase of the ML pipeline. In order to reveal these causal relationships, the Fast Greedy Search (FGES) algorithm was used. The algorithm was applied twice for each of the feature subsets obtained in the FS experiments. First, the algorithm was applied only to the features, excluding the outcome, and, then, it was applied to both the features and the outcome. The expectation was to find few or no differences between features that did not involve the outcome. That would mean that the CBNs are stable enough to draw some clear conclusions. The CBNs were built using the whole dataset with the selected features imputed by RF.

For the implementation and more detailed information about the methods, please refer to the corresponding sections.

## Dataset description

This study is part of the prospective observational trial *ShockOmics* European research project (“Multiscale approach to the identification of molecular biomarkers in acute heart failure induced by shock”, Nr. 602706 - ClinicalTrials.gov Identifier NCT02141607) [34] and was approved by: the Geneva Regional Research Ethics Committee (study number 14-041), the Ethical Committee of Hôpital Erasme-Université Libre de Bruxelles (study number P2014/171), and the Mútua de Terrassa Hospital Institutional Review Board (study number EO/1407). The main objective of this project was finding transcriptomics, proteomics and metabolomics signatures of shock as well as finding a robust set of features that could apply to all shock patients regardless of the initial injury. Patients admitted with both cardiogenic and septic shock to the intensive care units (ICU) of Geneva University Hospitals, Geneva, Switzerland (38-bed mixed) and Erasme Univesity Hospital, Brussels, Belgium (36-bed mixed) and Mutua de Terrassa (12-bed mixed) between October 2014 and March 2016 were screened for inclusion criteria. This study included patients diagnosed with septic or cardiogenic shock who were older than 18 years with a Sequential Organ Failure Assessment (SOFA) greater than 6, arterial lactate greater or equal than 2 mmol/l and with a documented source of infection at admission for patients with septic shock. Patients with a high risk of death within the first 24h after admission, systemic immunosuppresion, haematological diseases, metastasic cancer, pre-existing dialysis, decompensated cirrhosis or who had received more than 4 units of red blood cells or any fresh frozen plasma before screening were excluded from the study [34].

The complete database integrates blood samples and hemodynamic recordings from septic shock and cardiogenic shock patients, and from septic patients, obtained in the ICU [34]. Clinical data and blood samples were collected for analysis at: 1) T1 < 16 h from T0; 2) T2 = 48 h after T0; 3) T3 = day 7 or before discharge or before discontinuation of therapy in case of fatal outcome [35]. Since a lot of patients either died or were discharged during the study, the dataset has a lot of missing values at later time-steps (especially at T3). A short summary of the clinical data can be found in Table 1.

**Table 1.** Clinical data summary.

| Parameter | Cardiogenic shock | Septic shock | p-values |
| --- | --- | --- | --- |
| Number of patients | 25 | 50 | - |
| Gender (Males:Females) | 20:5 | 34:16 | - |
| Outcome (Alive:Dead) | 19:6 | 38:12 | - |
| Body mass index | $26.8 \pm 5.93$ | $25.22 \pm 5.54$ | 0.2591 |
| Total days in ICU | $6.6 \pm 5.42$ | $7.56 \pm 6.54$ | 0.5289 |
| SOFA<sub>T1</sub> | $10.72 \pm 4.27$ | $11.50 \pm 3.7$ | 0.4165 |
| SOFA<sub>T2</sub> | $8.33 \pm 4.92$ | $8.77 \pm 4.3$ | 0.699 |
| SOFA<sub>T3</sub> | $5.87 \pm 3.68$ | $6.75 \pm 4.7$ | 0.5334 |
| APACHE II<sub>T1</sub> | $24.12 \pm 9.79$ | $23.16 \pm 7.52$ | 0.6396 |
| APACHE II<sub>T2</sub> | $18.42 \pm 9.41$ | $16.36 \pm 8.45$ | 0.3530 |
| APACHE II<sub>T3</sub> | $12.8 \pm 5.35$ | $14.83 \pm 5.93$ | 0.2727 |
| Lactate levels<sub>T1</sub> | $6.04 \pm 4.3$ | $4.84 \pm 2.58$ | 0.1431 |
| Lactate levels<sub>T2</sub> | $1.52 \pm 0.91$ | $2.1 \pm 1.52$ | 0.1126 |
| Lactate levels<sub>T3</sub> | $1.25 \pm 0.5$ | $1.43 \pm 0.58$ | 0.3309 |

Summary of patient populations with cardiogenic and septic shock. Numerical parameters of both populations ( $mean \pm std$ ) are compared against each other with the Welch’s t-test, where the null hypothesis is that both parameters have identical mean values.

The *ShockOmics* dataset (data are available from the ShockOmics Institutional Data Access Board for researchers who meet the criteria for access to confidential data) has 333 features and 75 observations. 32.1% of all values are missing, and the highest percentage of missing values in one feature is 97%. Fig 1 shows a graphical depiction map of missing values for each observation. The dataset has both numerical and categorical values, but the majority of the data is numerical.

**Fig 1.**
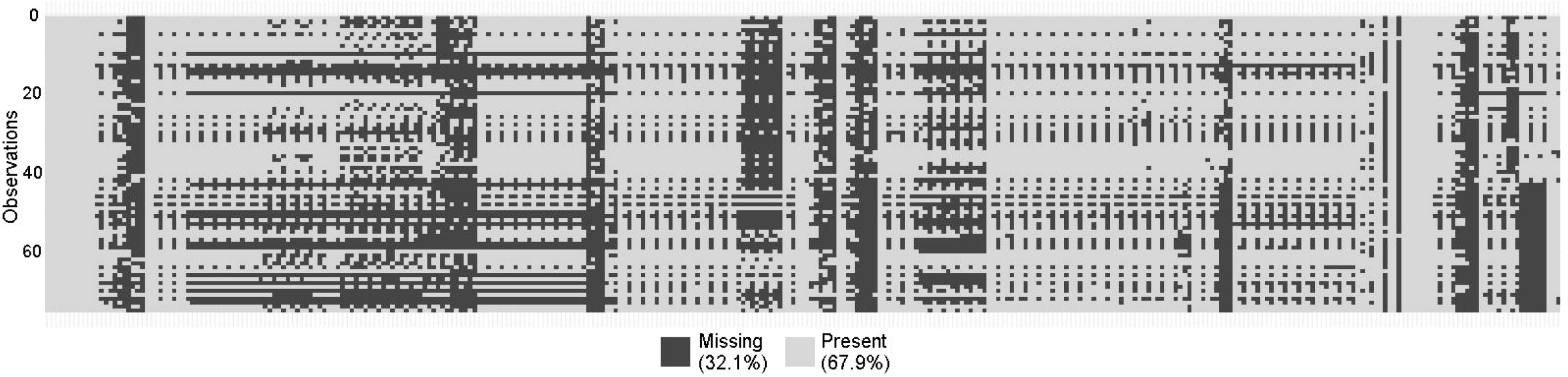
Graphical depiction map of missing values in *ShockOmics* dataset. The rows and the columns of the map correspond to observations and features, respectively. The black color represents missing values, the white color corresponds to present values in the *ShockOmics* dataset.

The *ShockOmics* dataset was split into three datasets. The first dataset was named *Full* and it was obtained by filtering features from the original *ShockOmics* dataset. Certain features that did not make sense from the classification viewpoint were manually removed. They were either very general comments in natural language or explicitly revealed some information about the outcome of the patient. Both cases were deemed not to be reliable as features to feed the ML model. In detail, the following features were manually removed: *ID*, *Reason for admission*; *ICU admission*; *RV area/LV area* (T1, T2, T3); *Microorganisms* (three columns with the same name); *ID*; *Death due to withdrawal of care*; *Mortality 28 days, 100 days* (two columns); *Hospital results*; *Total days in ICU, in Hospital* (two columns). Moreover, the features with more than 75% of missing data were also removed. The resulting dataset had 286 features with one target feature.

A research hypothesis is that the closer in time the feature to the final outcome, the better the prediction of mortality. To test this hypothesis, two further datasets were created, namely **T1+T2** and **T1**. Both were obtained from the *Full* dataset by filtering features at certain time-steps. From the *T1+T2* dataset all T3 features were removed, resulting in 206 features with a target feature. From the *T1* dataset both T2 and T3 time-step features were left out, so the resulting dataset included 114 features and a target feature.

A fourth feature set was built with features that are commonly assumed to be associated with the mortality of patients with shock, according to current practice. This feature set is referred to as *initial feature set* (*IFS*). Table 2 shows features that were included in the IFS set.

**Table 2.** Initial feature set description.

| Name of the feature | Observations | Type |
| --- | --- | --- |
| Which type of shock | 75 | categorical (4 val) |
| Lactate levels (mmol/L)<sub>T1</sub> | **72** | numerical (cont) |
| Lactate levels (mmol/L)<sub>T2</sub> | **64** | numerical (cont) |
| Lactate levels (mmol/L)<sub>T3</sub> | **38** | numerical (cont) |
| Mean arterial pressure (mmHg)<sub>T1</sub> | 75 | numerical (cont) |
| Mean arterial pressure (mmHg)<sub>T2</sub> | **71** | numerical (cont) |
| Mean arterial pressure (mmHg)<sub>T3</sub> | **44** | numerical (cont) |
| SOFA<sub>T1</sub> | 75 | numerical (cont) |
| SOFA<sub>T2</sub> | **71** | numerical (cont) |
| SOFA<sub>T3</sub> | **43** | numerical (cont) |
| APACHE II<sub>T1</sub> | 75 | numerical (cont) |
| APACHE II<sub>T2</sub> | 71 | numerical (cont) |
| APACHE II<sub>T3</sub> | 44 | numerical (cont) |
| Result in ICU | 75 | categorical (2 val) |

The columns of the table correspond to the name of the feature, the number of available observations and the type of the feature. *Initial feature set (IFS)* consists of 13 features and the target (*Result in ICU*). Features with missing observations are highlighted in bold in the *Observations* column.

## Feature selection and classification

In order to identify promising feature sets, five FS techniques were applied to the first three data sets described in the previous section. The resulting feature subsets were used to train ML models and five performance measures were recorded. The first four measures are defined through the numbers of true positives (TP), true negatives (TN), false positives (FP) and false negatives (FN):

- **Accuracy**: 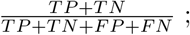
- **Matthews correlation coefficient (*MCC*)**: 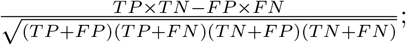
- **Sensitivity**: 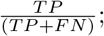
- **Specificity**: 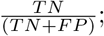
- **Area Under the Receiver Operating Characteristic Curve (*AUC*)**. The average AUC for both labels, weighted by the number of true instances for both labels.

The experiment pipeline for the FS procedure can be described as follows. First, 100 random splits were created for each of the datasets, using 75% for training and the rest for testing. Then, for each pair of training and test sets RF imputation [43] was used. Both sets were imputed separately one after another, using an imputed training set for a test set imputation. After that, the FS methods were applied, varying the size of the desired feature subset from a minimum of 2 to a maximum of 60. These FS models were subsequently trained on 75% of data, where only selected features were left in both training and test sets. The feature set that achieved the highest AUC was chosen as the best one for the split. The same procedure was applied to the rest of the 99 pairs of training and test sets. Once all feature sets and their performance measures were recorded, the mean and the standard deviation were calculated for each FS method and dataset. The complete experimental pipeline is graphically described in Figure 2.

**Fig 2.**
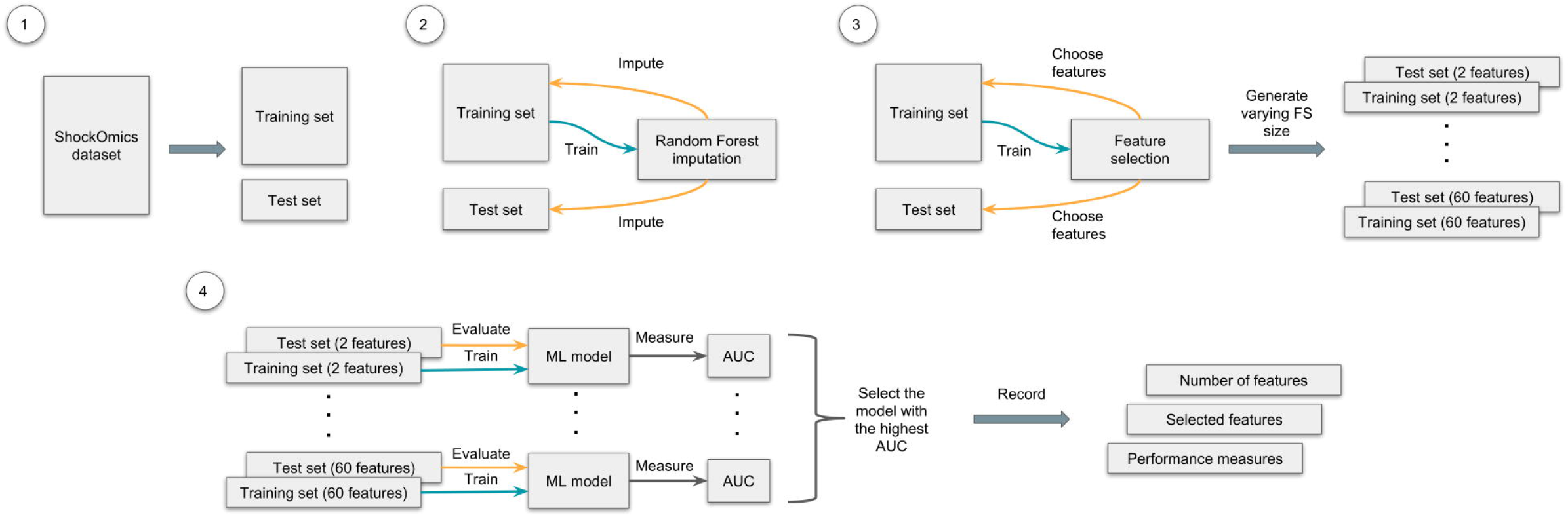
Graphical representation of the experimental pipeline. The process of FS and classification consists of the following steps: 1) create 100 random splits of the dataset: 75% for training and 25% for testing; 2) for each split impute both sets separately, using the imputed training set for a test set imputation; 3) after that, use the FS technique on the imputed training set, varying the size of the selected feature set (from a minimum of 2 to a maximum of 60), then choose the selected features in both sets, creating new pairs of training and test sets; 4) finally, using sets with different amounts of features, train and evaluate a ML model and choose the one with the highest AUC; record which and how many features were used for training this model, and its performance measures. Repeat these steps for all 100 random splits for each dataset and FS technique.

Now that the best feature sets in all pairs of splits were selected, the stability scores were calculated as frequencies: the individual feature occurrences in all chosen sets were counted and then divided by the number of pairs (100 in the reported experiments). In the S1 Appendix, the 80 features with the highest stability scores are listed for each FS technique and dataset, but note that only the best 20 were later chosen to build CBNs.

The RF data imputation was performed in *R* (v3.4.3) [41], using *missForest* package that allows to impute continuous and/or categorical data including complex interactions and nonlinear relations.

The scikit-learn [36] library (v0.19.1) in Python (v2.7) was used for the implementation of the FS and ML methods. The default parameters of the methods were used in the experiments, if not otherwise specified. The five FS techniques used in the identification of promising attributes and the corresponding ML models for feature evaluation are described next:

- **Univariate feature selection based on ANOVA F-values (*UFS*)**. Univariate feature selection [37] works by selecting the best features based on univariate statistical tests. In this case, ANOVA F-tests are used. The Gaussian Naive Bayes model [42] was used for the evaluation. The UFS was implemented using the sklearn.feature selection.SelectKBest method that removes all but the *k* highest scoring features and sklearn.feature selection.f classif for the ANOVA F-tests. The Naive Bayes model was implemented through the sklearn.naive bayes.GaussianNB method.
- **Recursive feature elimination with SVC (*RFE*)**. Given an external estimator that assigns weights to features (e.g., the coefficients of a linear model), the goal of the RFE is to select features by recursively considering smaller and smaller sets of features [38]. The RFE was implemented through the sklearn.feature selection.RFE method. For the external estimator and, as the evaluated ML model, the SVC [39] was used with the help of the sklearn.svm.SVC method;
- **Two stage FS (*UFS + RFE*)**. A combination of two FS techniques from this list. In the first stage, UFS preselects the 120 best features (80 for the T1 dataset) based on the ANOVA F-value and then RFE (SVC) chooses the best features in the second stage. Here, the SVC was used as the ML model for the evaluation;
- **Random Forest (*RF*) FS**. This method [38] is based on the trained RF model [40] and its feature rankings. The sklearn.feature selection.SelectFromModel method was used to retrieve the feature rankings and the sklearn.ensemble.RandomForestClassifier method to train the RF model for the FS and later (with only selected features) for evaluation (10 trees were used in each stage);
- **Aggregated (*Aggr*.) FS**. The method uses the averaged stability scores for FS obtained by the four previous techniques and, therefore, does not include the ML model. Instead, the performance is measured by averaging the measures obtained by the previous methods. This FS does not involve any processing of the dataset, but relies heavily on the previous FS techniques. As a result, it allows to achieve more stable features and analyze the aggregated performance of all trained ML models for the dataset.

## Structure learning

The FGES algorithm was used to obtain the CBN structures that would reveal the interactions between the features. This is an optimized and parallelized version of the Greedy Equivalence Search algorithm. It heuristically searches the space of CBNs and returns the model with the highest value for the Bayesian Information Criterion (BIC) [45] it finds. It starts with an empty graph and adds new edges to increase the Bayesian score. The process stops when there are no edges that increase the score. It then starts removing edges until no single edge removal can increase the score.

The algorithm works on the assumption that the causal process generating the data is accurately modeled by a CBN. Each node in this CBN is a a linear function of its parents, plus a finite additive Gaussian noise term. Each observation in the data is assumed to be independent and obtained by randomly sampling all the variables from the joint distribution. Given all these assumptions, the FGES procedure outputs the CBN structure that contains:

1. an arc X → Y, if and only if X causes Y;
2. an edge (-), if and only if either X causes Y or Y causes X;
3. no edge between X and Y, if and only if X and Y have no direct causal relationship between them.

The *Tetrad* software (v6.4.0) was used in the implementation of FGES. The algorithm was applied twice for each of the feature sets obtained during the feature selection phase. First, the algorithm was applied only to the features, excluding the outcome, while the second time it was instead applied to both features and the outcome. The expectation was to have few or no differences between features that did not involve the outcome. That would mean that the CBNs are stable enough to draw clear conclusions. The algorithm used the default parameters and seven maximum discrete categories.

## Experimental Results

### Feature selection experiments

The FS experiments are divided into three groups. Each group corresponds to its own dataset: *Full*, *T1+T2* and *T1*. All five FS techniques were applied to each dataset. The results were used to train the ML model, and the performance of each feature set was evaluated for each model. In this section, the best models from all FS experiments and their accuracy, MCC, sensitivity, specificity and AUC are presented. For the details on individual features and their stability scores please refer to the S1 Appendix. Considering the 80 features with the highest stability scores for each FS technique and dataset, one may appreciate how the selected features vary across the 100 random splits.

Table 3 shows the comparison of different FS techniques and the general performance of the best features they chose: the size of the feature set, the dataset and the FS method used. The FS techniques that were applied to the same dataset are grouped together. The additional IFS dataset is used as a baseline to compare the rest of feature sets performances (the Gaussian Naive Bayes was chosen as the ML model). A naive *majority classifier* that always predicts the most frequent label in the training set, was also added for baseline comparison.

**Table 3.** The performance comparison of the feature sets.

| # Features | Data | Method | Accuracy | MCC | Sensitivity | Specificity | AUC |
| --- | --- | --- | --- | --- | --- | --- | --- |
| - | - | Majority class | $0.763 \pm 0.09$<br>(0.0) | $0.0 \pm 0.0$<br>(0.0) | $0.0 \pm 0.0$<br>(0.0) | $1.0 \pm 0.0$<br>(0.0) | $0.5 \pm 0.0$<br>(0.0) |
| 11 | IFS | - | $0.644 \pm 0.124$<br>(0.0) | $0.028 \pm 0.255$<br>(0.0) | $0.269 \pm 0.252$<br>(0.0) | $0.767 \pm 0.14$<br>(0.0) | $0.518 \pm 0.139$<br>(0.0) |
| $13.5 \pm 13.6$ | T1 | UFS | $0.828 \pm 0.083$<br>(0.0) | $0.573 \pm 0.183$<br>(0.0) | $0.731 \pm 0.217$<br>(0.2496) | $0.873 \pm 0.095$<br>(0.0) | $0.802 \pm 0.108$<br>(0.0037) |
| $16.5 \pm 13.2$ | T1 | RFE | $0.823 \pm 0.079$<br>(0.0) | $0.549 \pm 0.19$<br>(0.0) | $0.702 \pm 0.23$<br>(0.0396) | $0.876 \pm 0.089$<br>(0.0001) | $0.789 \pm 0.114$<br>(0.0003) |
| $14.5 \pm 13.0$ | T1 | UFS+RFE | $0.814 \pm 0.077$<br>(0.0) | $0.52 \pm 0.181$<br>(0.0) | $0.656 \pm 0.226$<br>(0.0004) | $0.876 \pm 0.092$<br>(0.0001) | $0.766 \pm 0.106$<br>(0.0) |
| $18.1 \pm 10.8$ | T1 | RF | **$0.87 \pm 0.07$**<br>(0.4397) | **$0.668 \pm 0.152$**<br>(0.268) | **$0.755 \pm 0.198$**<br>(0.7221) | **$0.922 \pm 0.074$**<br>(0.7848) | **$0.839 \pm 0.092$**<br>(0.6576) |
| $15.7 \pm 12.8$ | T1 | Aggr. | $0.834 \pm 0.08$<br>(0.0001) | $0.577 \pm 0.186$<br>(0.0) | $0.711 \pm 0.221$<br>(0.0709) | $0.887 \pm 0.09$<br>(0.002) | $0.799 \pm 0.109$<br>(0.0021) |
| $10.8 \pm 14.2$ | T1+T2 | UFS | $0.839 \pm 0.085$<br>(0.0008) | $0.584 \pm 0.207$<br>(0.0001) | $0.715 \pm 0.229$<br>(0.101) | $0.889 \pm 0.091$<br>(0.0035) | $0.802 \pm 0.116$<br>(0.0053) |
| $17.5 \pm 15.8$ | T1+T2 | RFE | $0.807 \pm 0.089$<br>(0.0) | $0.503 \pm 0.205$<br>(0.0) | $0.637 \pm 0.265$<br>(0.0002) | $0.874 \pm 0.101$<br>(0.0001) | $0.755 \pm 0.122$<br>(0.0) |
| $12.9 \pm 10.7$ | T1+T2 | UFS+RFE | $0.796 \pm 0.074$<br>(0.0) | $0.465 \pm 0.175$<br>(0.0) | $0.596 \pm 0.251$<br>(0.0) | $0.873 \pm 0.097$<br>(0.0001) | $0.734 \pm 0.109$<br>(0.0) |
| $21.2 \pm 14.4$ | T1+T2 | RF | **$0.878 \pm 0.076$**<br>(1.0) | **$0.693 \pm 0.166$**<br>(1.0) | **$0.765 \pm 0.199$**<br>(1.0) | **$0.925 \pm 0.081$**<br>(1.0) | **$0.845 \pm 0.099$**<br>(1.0) |
| $15.6 \pm 13.9$ | T1+T2 | Aggr. | $0.83 \pm 0.087$<br>(0.0) | $0.561 \pm 0.208$<br>(0.0) | $0.678 \pm 0.246$<br>(0.0065) | $0.89 \pm 0.095$<br>(0.0056) | $0.784 \pm 0.12$<br>(0.0001) |
| $17.9 \pm 17.2$ | Full | UFS | $0.754 \pm 0.105$<br>(0.0) | $0.31 \pm 0.23$<br>(0.0) | $0.431 \pm 0.256$<br>(0.0) | $0.861 \pm 0.129$<br>(0.0) | $0.646 \pm 0.119$<br>(0.0) |
| $19.7 \pm 16.5$ | Full | RFE | $0.758 \pm 0.088$<br>(0.0) | $0.334 \pm 0.213$<br>(0.0) | $0.483 \pm 0.278$<br>(0.0) | $0.855 \pm 0.114$<br>(0.0) | $0.669 \pm 0.118$<br>(0.0) |
| $17.4 \pm 15.2$ | Full | UFS+RFE | $0.743 \pm 0.09$<br>(0.0) | $0.311 \pm 0.21$<br>(0.0) | $0.477 \pm 0.251$<br>(0.0) | $0.839 \pm 0.123$<br>(0.0) | $0.658 \pm 0.111$<br>(0.0) |
| $21.9 \pm 15.5$ | Full | RF | **$0.819 \pm 0.082$**<br>(0.0) | **$0.53 \pm 0.176$**<br>(0.0) | **$0.614 \pm 0.244$**<br>(0.0) | **$0.902 \pm 0.087$**<br>(0.0544) | **$0.758 \pm 0.111$**<br>(0.0) |
| $19.2 \pm 16.2$ | Full | Aggr. | $0.769 \pm 0.096$<br>(0.0) | $0.371 \pm 0.228$<br>(0.0) | $0.501 \pm 0.267$<br>(0.0) | $0.864 \pm 0.117$<br>(0.0) | $0.683 \pm 0.123$<br>(0.0) |

Best feature sets obtained in the FS experiments and their general performance across 100 random data splits. The columns correspond to the number of features, the data that were used to obtain the feature set, the FS method and five performance measures ( $mean \pm std(p - value)$ ). Welch's t-test was used to obtain p-values for the null hypothesis that two performance measures had identical values. Each measure was tested against the same measures of the (T1+T2, RF) model. The *IFS* dataset was used as a baseline for comparison. The best results for all measures in each dataset group are highlighted in bold.

As it can be seen from Table 3, the FS models outperform manually chosen features from the IFS. Moreover, the IFS model is only slightly better than the naive *majority classifier*. The data is unbalanced and that is the reason why the accuracy and the specificity are the only measures for which the *majority classifier* is better. As for the rest of the models, they show consistently better results than the baseline. The only exception is the specificity measure, which means that there were no models that identified all negative examples correctly across all 100 random splits, and the accuracy for the *RFE* and *UFS+RFE* techniques with the *Full* dataset. The *RF* models produced consistently good results in all datasets, and were considered best for every dataset. The best result of all experiments were shown by one of these models for the *T1+T2* dataset. Based on the *Aggr*. performance, the general performance of the models declined with the additional time-steps. This can be explained by the lower performance of the *RFE* and *UFS+RFE* models for the *T1+T2* dataset, compared to the *T1*, and by the fact that the *Full* models performed the worst across all three datasets.

For the causal discovery experiments, it was decided to choose the 20 features with the highest stability scores for the models whose results are shown in Table 4. The *RF* models showed the best performance and therefore their features required more thorough investigation. All feature sets selected for the causal discovery experiments and their individual features can be found in the list below.

**Table 4.** The performance of the feature sets selected for the causal discovery experiments.

| # Features | Data | Method | Accuracy | MCC | Sensitivity | Specificity | AUC |
| --- | --- | --- | --- | --- | --- | --- | --- |
| $18.1 \pm 10.8$ | T1 | RF | $0.87 \pm 0.07$<br>(0.4397) | $0.668 \pm 0.152$<br>(0.268) | $0.755 \pm 0.198$<br>(0.7221) | $0.922 \pm 0.074$<br>(0.7848) | $0.839 \pm 0.092$<br>(0.6576) |
| $15.7 \pm 12.8$ | T1 | Aggr. | $0.834 \pm 0.08$<br>(0.0001) | $0.577 \pm 0.186$<br>(0.0) | $0.711 \pm 0.221$<br>(0.0709) | $0.887 \pm 0.09$<br>(0.002) | $0.799 \pm 0.109$<br>(0.0021) |
| $21.2 \pm 14.4$ | T1+T2 | RF | **$0.878 \pm 0.076$**<br>(1.0) | **$0.693 \pm 0.166$**<br>(1.0) | **$0.765 \pm 0.199$**<br>(1.0) | **$0.925 \pm 0.081$**<br>(1.0) | **$0.845 \pm 0.099$**<br>(1.0) |
| $15.6 \pm 13.9$ | T1+T2 | Aggr. | $0.83 \pm 0.087$<br>(0.0) | $0.561 \pm 0.208$<br>(0.0) | $0.678 \pm 0.246$<br>(0.0065) | $0.89 \pm 0.095$<br>(0.0056) | $0.784 \pm 0.12$<br>(0.0001) |
| $21.9 \pm 15.5$ | Full | RF | $0.819 \pm 0.082$<br>(0.0) | $0.53 \pm 0.176$<br>(0.0) | $0.614 \pm 0.244$<br>(0.0) | $0.902 \pm 0.087$<br>(0.0544) | $0.758 \pm 0.111$<br>(0.0) |
| $19.2 \pm 16.2$ | Full | Aggr. | $0.769 \pm 0.096$<br>(0.0) | $0.371 \pm 0.228$<br>(0.0) | $0.501 \pm 0.267$<br>(0.0) | $0.864 \pm 0.117$<br>(0.0) | $0.683 \pm 0.123$<br>(0.0) |

The columns of the table correspond to the number of features, the data that was used to obtain the feature set, the FS method used and five performance measures (*mean ± std(p value)*). Each measure was tested against the same measures of the ( $T1+T2$ , *RF*) model. The best results in the performance measures columns are highlighted in bold.

[T1, RF]: APACHE II_T1, SOFA_T1, Respiratory rate_T1, Urine Output (mL/ day)_T1, E wave (cm/s)_T1, Fluid Balance (ml)_T1, K Ur_T1, Lactate levels (mmol/L)_T1, Tidal volume (VT)_T1, Pulmonary artery systolic pressure (TR jet by CW + CVP) (mmHg)_T1, * Norepinephrine (mg/kg/min)_T1, PCT_Value (mg/mL)_T1, Heart rate (bpm)_T1, E/e’_T1, PEEP_T1, Pplat_T1, Platelet count_T1, Na Ur_T1, pH_T1, PaO2/FiO2_T1;

[T1, Aggr.]: APACHE II_T1, SOFA_T1, Respiratory rate_T1, Glasgow Coma Scale_T1, K Ur_T1, Platelet count_T1, Tidal volume (VT)_T1, Tricuspid regurgitation maximal velocity (by CW) (cm/s)_T1, HCO3 (mmol/L)_T1, LA dilatation by eyeballing_T1, Pulmonary artery systolic pressure (TR jet by CW + CVP) (mmHg)_T1, Base Excess (mmol/L)_T1, E wave (cm/s)_T1, PCT_Value (mg/mL)_T1, Respiratory rate (rpm)_T1, Pplat_T1, Tricuspid annular tissular doppler S wave (DTI) (cm/s)_T1, * Norepinephrine (mg/kg/min)_T1, * Dobutamine (mg/kg/min)_T1, Lactate levels (mmol/L)_T1;

[T1+T2, RF]: Lactate levels (mmol/L)_T2, SOFA_T2, Tidal volume (VT)_T2, APACHE II_T1, Urine Output (mL/day)_T1, Urine Output (mL/day)_T2, APACHE II_T2, Respiratory rate_T1, K Ur_T2, SOFA_T1, * Norepinephrine (mg/kg/min)_T2, Neutro abs count_T2, E wave (cm/s)_T1, Na Ur_T2, pH_T2, K Ur_T1, Glasgow Coma Scale_T2, Tricuspid regurgitation maximal velocity (by CW) (cm/s)_T1, PT_T2, Pulmonary artery systolic pressure (TR jet by CW + CVP) (mmHg)_T1;

[T1+T2, Aggr]: APACHE II_T1, SOFA_T2, Tricuspid regurgitation maximal velocity (by CW) (cm/s)_T1, Glasgow Coma Scale_T1, Neutro abs count_T2, Glasgow Coma Scale_T2, Platelet count_T1, K Ur_T1, APACHE II_T2, SOFA_T1, Respiratory rate_T1, HCO3 (mmol/L)_T1, Lactate levels (mmol/L)_T2, E wave (cm/s)_T1, Tidal volume (VT)_T1, Respiratory rate (rpm)_T1, K Ur_T2, Na Ur_T2, PT_T2, LA dilatation by eyeballing_T1;

[Full, RF]: Neutro abs count_T1, Platelets (10ˆ3/mmˆ3)_T3, E/e’_T2, Lateral e’ (cm/s)_T2, Creat Ur_T2, Urine Output (mL/day)_T2, Lateral e’ (cm/s)_T1, Respiratory rate (rpm)_T1, Lympho abs count_T1, Mean arterial pressure (mmHg)_T1, Diastolic Blood Pressure (mmHg)_T2, E wave deceleration time (ms)_T2, Tricuspid annular tissular doppler S wave (DTI) (cm/s)_T1, Sat O2/FiO2_T2, Platelets (10ˆ3/mmˆ3)_T1, FiO2_T3, Tidal volume (VT)_T1, RBC count_T3, Fluid Balance (ml)_T2, Platelet count_T2;

[Full, Aggr]: A wave (cm/s)_T2, Systolic blood pressure (mmHg)_T2, Heart rate (bpm)_T1, Respiratory rate (rpm)_T1, Weight (kg), Height (cm), A wave (cm/s)_T1, PT_T1, E wave deceleration time (ms)_T3, Hematocrit (%)_T2, Platelet count_T3, Creat Ur_T1, E wave deceleration time (ms)_T1, PaCO2 (mmHg)_T1, K Ur_T1, E wave (cm/s) _T2, aPTT_T1, Mean arterial pressure (mmHg)_T1, Base Excess (mmol/L) _T1, PaO2 (mmHg)_T2.

List of best feature selections. Within square brackets: the subset of data that was used to obtain the feature set and the applied FS technique.

## Causal discovery

For each of the feature sets there are two corresponding CBNs. The first one (the CBN ’a’ was built without the target feature (*Result in ICU*), the second one (the CBN ’b’) included all features and the target. This section contains the CBNs and their descriptions for the (*T1, RF*) and (*T1+T2, RF*) feature sets. For the rest of CBNs, please refer to the S1 Appendix.

Fig 3 shows the CBNs for the (*T1, RF*) feature set. As it can be seen, two nodes, namely *E wave* and *E/e* at T1 are detached from the rest of the network in both CBNs. In CBN ’a’, all edges are undirected, but the addition of the target feature, *Result in ICU*, changes most of them to directed. Moreover, adding this feature replaces the edge between *Lactate levels* and *Norepinephrine* at T1 to the edge between *K Ur* and *Na Ur* at T1, connecting these two nodes to the CBN. According to the CBN ’b’, *Result in ICU* can be potentially influenced by *Norepinephrine*, *Respiratory rate*, *Platelet count* and *SOFA* at T1.

**Fig 3.**
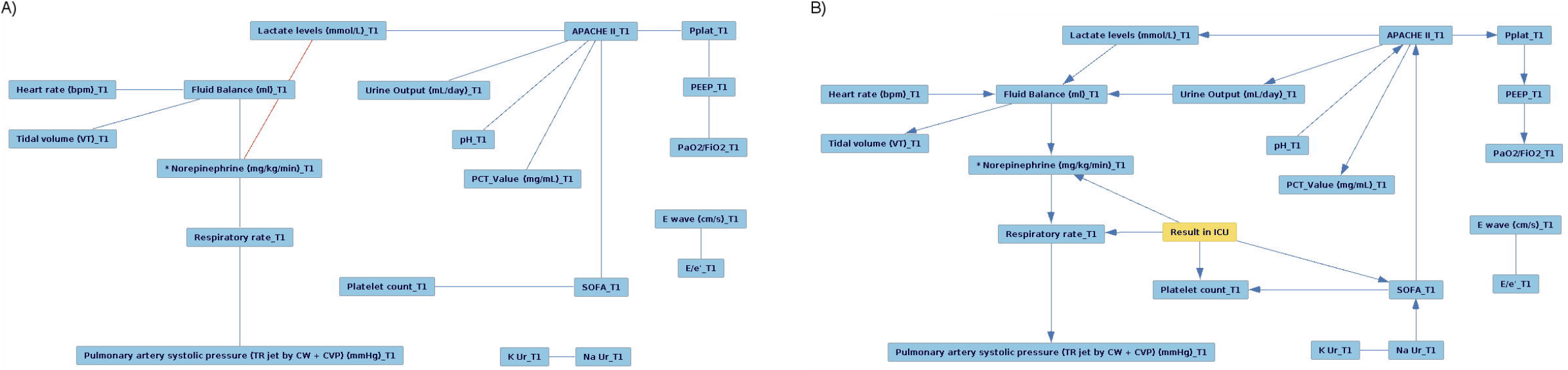
The (*T1, RF*) CBN. The CBN for the (*T1, RF*) feature set, obtained: a) without the target feature; b) with the target feature.

Fig 4 shows the CBNs for the (*T1+T2, RF*) feature set. In both CBNs, no edges are connected to *K Ur* at T1. This time, the addtion of the outcome attaches *E wave* at T1 and *Neutro abs count* at T2 to the rest of the network. There are multiple changes across the CBN that involve other features as well. For example, edges between *Glasgow Coma Scale* at T1 and *Lactate levels* at T2, and between *Lactate levels* at T2 and *PT* at T2 disappear. Moreover, for all edges the causal directions become clear, and new edges emerge. The CBN ’b’ shows that *Result in ICU* is potentially influenced by *E wave* at T1, *Norepinephrine* at T2, *Neutro abs count* at T2 and *SOFA* at T2.

**Fig 4.**
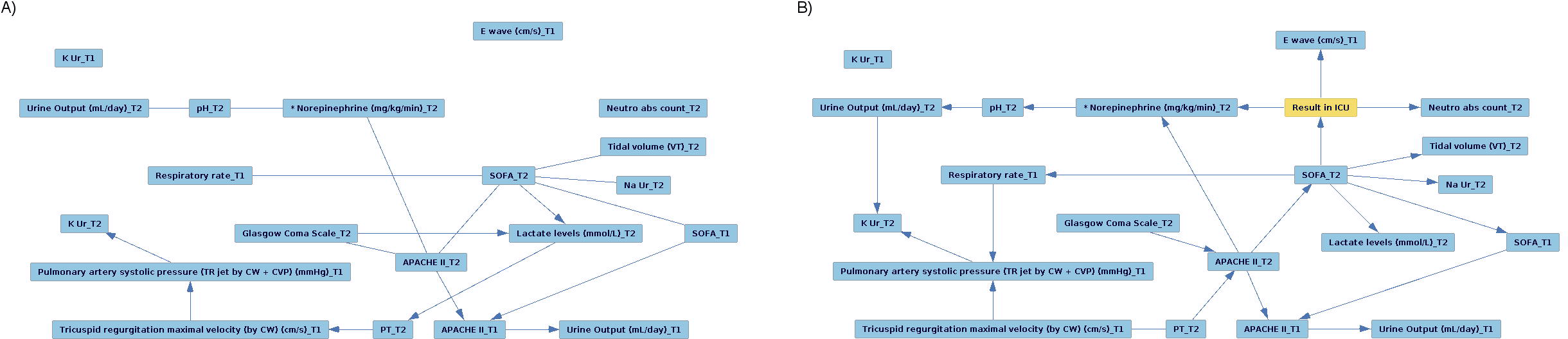
The (*T1+T2, RF*) CBN. The CBN for the (*T1+T2, RF*) feature set obtained: a) without the target feature; b) with the target feature.

In most of CBNs pairs, the presence of the target value did not imply much change. The majority of edges of the *Full* models were undirected and the parts of the network were detached from each other. Adding *Result in ICU* does not connect these separate parts. This contrasts with the *T1* and *T1+T2* models, that have more or less connected CBNs. As for the *T1* and T1+T2 CBNs, they have common patterns that are noticeable throughout the experiments. More importantly, the two best models described in this section have some common features that are close to the outcome in both CBNs. These similarities are discussed in the next section.

## Discussion

Analyzing the results of the FS experiments, one may notice that there are a lot of reoccurring features with high stability scores for each of the datasets. These features can be clearly seen from the *Aggr*. tables in page 19. For *T1* the most common features are *APACHE II*, *SOFA*, *respiratory rate*, *Glasgow coma scale* and *K Ur*. All these features except *Glasgow coma scale* scored high with the model that showed the best results for this dataset. Moreover, some of them were also present in the results for *T1+T2*. Interestingly, there are many similar features that scored high in both *T1* and *T1+T2*, which is a clear indication of their importance for the mortality prediction task.

Regarding the *T1+T2* dataset, the most stable features are *APACHE II* at T1, *SOFA* at T2, *tricuspid regurgitation maximal velocity* at T1, *Glasgow Coma Scale* at T1 and *neutrophils absolute count* at T2. It is especially interesting that some attributes at T1 occurred more frequently than their T2 counterparts. This can be explained by the fact that features at later time-steps present more noise and missing features. However, the (*T1+T2, RF*) showed the best predictive power in our experiments, which means that at least some T2 features improved the results obtained.

In contrast, the worst performance was that obtained for the *Full* dataset. Its most stable features include *A wave* at T2, *systolic blood pressure* at T2, *heart rate* at T1, *respiratory rate* at T1 and *weight*. These features scored the highest, but their stability scores are very low. Moreover, the difference between all scores of the *Full* dataset is very small. This means that during the FS procedures across 100 random experiments, very different features were selected making them unreliable for their low stability. This supports the hypothesis that adding more time-steps (i.e., later times) increases noise, and that the current amount of data is not sufficient enough to bypass this problem.

The performance of features from different datasets and models were further compared to the model that showed the best results, which was the (*T1+T2, RF*) model. More specifically, the performance measures of (*T1+T2, RF*) were tested against the measures of other models, using the Welch’s t-test. The null hypothesis was that each pair of performance measures has identical values. Setting the threshold for the p-value to 0.05 shows that there is a significant chance that the (*T1, RF*) and (*T1+T2, RF*) models perform similarly according to all five performance measures. For this reason, these models were singled out in the causal discovery experiments.

The CBN structures revealed connections between the features and the outcome. The assumption behind the evaluation of the most important clinical attributes for outcome prediction was that such features are the closest to the outcome in a graph. The direction of the edge from feature to the outcome is also a good indicator that a certain feature is important.

According to the *RF* and *Aggr*. CBNs for the *T1* dataset, *Result in ICU* is influenced by *norepinephrine* and *SOFA* at T1. Both models presented direct connections between these two features and the outcome. The additional possible candidates, supported by the (*T1, RF*), are *respiratory rate* and *platelet count* at T1. *E wave*, *dobutamine* and *tricuspod regurgitation maximal velocity* at T1 were considered important by the (*T1, Aggr*.) pair. As for the *T1+T2* dataset, its *RF* and *Aggr*. models show the significance of *neutrophils absolute count* at T2. Additional features that are directly connected to *Result in ICU* in the *RF* CBN include *norepinephrine* at T2, *E wave* at T1 and *SOFA* at T2. The *Aggr*. model adds *APACHE II* at T2 to the list. The *Full* CBNs show that the selected features do not create any coherent structure and, hence, cannot be used to draw any conclusions. This can be explained by the low stability scores and the lack of variability among them in the previous experiments.

Some parallels can be seen between the features that achieved high *Aggr*. stability scores and those that were considered important in the causal discovery experiments. These features were ranked very high by the FS techniques and had the direct connection to the outcome. These features include *SOFA* at T1, *respiratory rate* at T1, *tricuspid regurgitation maximal velocity* at T1, *neutrophils absolute count* at T2 and *SOFA* at T2. Additional features that were in the top 20 of the *Aggr*. stability scores tables but were ranked lower are *norepinephrine* at T1, *platelet count* at T1, *E wave* at T1 and *dobutamine* at T1. The full list of features that were considered valuable for the mortality prediction task can be found below. Since the *Full* features had very low stability scores, only the first few T1 and T1+T2 features from *Aggr*. tables were considered important.

High stability scores features: APACHE II T1, SOFA T1, Respiratory rate T1, Glasgow Coma Scale T1, K Ur T1, SOFA T2, Tricuspid regurgitation maximal velocity T1, Neutro abs count T2, Glasgow Coma Scale T2.

Causal discovery features: Norepinephrine T1, SOFA T1, Respiratory rate T1, Platelet count T1, E wave at T1, Dobutamine T1, Tricuspod regurgitation maximal velocity T1, Neutro abs count T2, Norepinephrine T2, SOFA T2, APACHE II T2.

List of attributes that were considered promising for the outcome prediction in patients with shock.

These findings agree with other studies that support the use of certain attributes for mortality prediction. For instance, the SOFA score has previously been shown to have a significant prognostic value for in-hospital mortality prediction [46] [47]. The APACHE II score is often used for outcome prediction too [48–50]. And in the study by Jansen *et al*. [51], targeting a decrease of at least 20% in the blood lactate level (a common feature for the best two models) over a 2-hour period seemed to be associated with reduced in-hospital mortality in patients with shock. However, some of the attributes that are also related to mortality in patients with shock, like mean blood pressure [52], mean arterial pressure [53], or low LVEF [54] did not show significant predictive value in this study.

Finally, it should be noted that some features seem more important at certain time-steps. For example, the difference in stability scores shows that *neutrophils absolute count* at T2, *tricuspid regurgitation maximal velocity* at T1 and *dobutamine* at T2 were selected far more often than their counterparts. This might be due to the specifics of imputation or due to the lack of data. In any case, this hypothesis requires a more thorough investigation, that is beyond the scope of this work.

## Conclusion

In this paper, we have analyzed and experimentally compared several models for the prediction of mortality in patients with cardiogenic or septic shock, identifying those clinical traits that are more relevant for such prediction in the ICU at different time-points. These models are the result of the application of different FS techniques over the available clinical data. In particular, we were interested in obtaining an actionable classifier for the acute phase of shock, which is the most critical to implement an appropriate therapeutic response and, therefore, has the greater impact on the prognosis of patients. For the acute phase (i.e. the first 24 hours of evolution of shock), we have obtained a classifier that leverages upon well established attributes for septic shock such as the SOFA or APACHE II scores and treatment with vasopressors (for example, norepinephrine), which are part of the cardiovascular SOFA score. This classifier for the acute phase of shock also resorts to other valuable attributes such as those obtained through echocardiography during cardiogenic shock (e.g. E wave velocity, early mitral inflow velocity and mitral annular early diastolic velocity (E/e’)). Other important attributes are related to respiratory function (such as respiratory rate, tidal volume and ventilator pressures), fluid balance, pulmonary artery pressure, renal function (such as urine output and urea levels), lactate levels, acidosis and C-reactive protein levels. These significant attributes are very closely related to the new official definition of shock and its corresponding management guidelines, in particular those related to organ dysfunction and respiratory function. However, our causal discovery experiments showed that treatment with pressors, respiratory rate, platelet count and the SOFA score presented the highest dependence with ICU outcome. The best classifier for the acute phase yielded an accuracy of 0.870, a sensitivity of 0.755 and a specificity of 0.922. The accuracy results were not statistically different from those obtained with the best classifier at T1 and T2.

The main conclusion of this study is that it is possible to predict the risk of death in the acute phase of septic and cardiogenic shock with quite acceptable results by taking into consideration the attributes routinely measured through echocardiography in the ICU. Use of this data for assessing the prognosis of patients is considered valuable for the clinical management of patients with shock in the ICU at ICU admission or during the first day of evolution.

The results presented here open up a new line of research for the study of the pathophysiology of shock when combined with other data related to organ dysfunction beyond the commonly used scales such as the SOFA score. In the next step of this research, we plan to add other types of data that are related to the patient’s response to shock and assess to what extend the expanded models may yield improved results. This new layer of data will include transcriptomics, proteomics and metabolomics features as detailed in the scope of the ShockOmics European project.

## Acknowledgments

This research was supported by the European FP7 project Shockomics (Nr. 602706), Multiscale approach to the identification of molecular biomarkers in acute heart failure induced by shock, and Spanish research project TIN2016-79576-R.

## Supporting information

S1 Appendix. Complementary material for the FS and causal discovery experiments. The resulting features from applying multiple FS techniques to the three available datasets, their stability scores, additional CBNs and a full list of ShockOmics attributes.

